# Complex modulation of visual attention to 3^rd^ person interactions in wild macaques

**DOI:** 10.1101/2025.09.23.677964

**Authors:** Sofia M. Pereira, Suthirote Meesawat, Suchinda Malaivijitnond, Julia Ostner, Oliver Schülke

## Abstract

Social animals rely on socio-cognitive skills to monitor their social environment and make informed decisions. Yet, visual attention to others’ interactions in real-life scenes remains understudied, despite evidence for dedicated neuronal networks for the processing of real-life social information and for social interactions in particular. Here, we ask how subject characteristics, interaction valence, and social relationships between subject and stimuli interact to guide overt attention of 56 wild male Assamese macaques to 962 3^rd^ person social scenes that unfolded spontaneously in their vicinity. Social interactions drew more and longer attention than social control scenes lacking the interaction, highlighting the relevance of interaction. The effect of both social bond strength and dominance relations between subject and stimulus on the probability to attend was independently modulated by scene valence. Moreover, beyond reacting to the affordances of the social scene and independent of their social status, males differed in how long they observed the interactions of others. These findings suggest that visual attention is guided by the value an individual associates with different aspects of the social scene based on previous experience in both the agonistic and the affiliative realms and by current threat indicative of complex information integration in the brain.

## INTRODUCTION

Social neuroscience research employs a variety of social stimuli that vary in their naturalism^1^. On one end of the spectrum the subject is chair-seated and presented with static images of social agents and non-social controls, while their eye gaze and neuronal activity is tracked^2–4^. From there, naturalism is increased by using videos that are more salient, but also noisier, stimuli^5,6^. Virtual reality scenes can even more closely mimic real-life interactions, as avatars can display contingent behavior creating a live-stream of interactions^7,8^. However, real-life agents are the most naturalistic, yet also the most challenging to control^9,10^ particularly in natural environments^1^. This study leverages the strengths of naturalistic animal observations, where large numbers of subjects can be studied allowing us to weigh individual differences in social attention against the affordances of the stimulus situation including subject’s affiliative and dominance relations with the stimuli.

Social attention, i.e. monitoring conspecifics to extract information, is a crucial building block of advanced social cognition^11^, involving specialized brain networks^12,13^. Social primates are willing to forgo an immediate food reward for the privilege to watch conspecifics in videos^14^. Yet, not all social information is equally relevant. For example, rhesus monkeys (*Macaca mulatta*) pay with forgone juice rewards for viewing a picture of the face of a dominant male, but have to be paid to look at a subordinate^15^. Visual attention for faces provides information about others’ overt spatial attention, i.e. about gaze direction^16^, and social signals encoded in ritualized facial expressions^2^. In fact, macaques pay more attention to facial expressions associated with threat than to those associated with benign intent and even less to neutral faces ^17,18^, and more readily follow the gaze of others if they show an emotional as opposed to a neutral expression ^19^. Single neurons in the amygdala show selective activity for threat, friendly and neutral facial expressions^2^.

Thus, social attention targets faces, gazes, and facial expressions as well as relevancies unrelated to perceptual aspects of the stimulus, e.g. stimulus dominance status, all of which can be tested with static information. Testing with video recorded dynamic stimuli shows that sustained attention depends on additional ethogram elements contained in the video sequence^20,21^: Rhesus macaques looked longer at areas of a frame with a grooming solicitation, aggression or submission^20^, and chimpanzees (*Pan troglodytes*) pay more attention to interacting dyads than solitary controls^21^. During fMRI, macaques engage a prefrontal cortical area specifically when watching videos of two others interacting compared to two others acting each on their own or simply resting; there is a brain for interaction^6^.

Presenting videos of longer natural scenes with multiple real-life agents hugely advances our understanding of selective social attention in complex environments and the associated neurophysiology^1,13^. However, dual eye-tracking and electro-physiology demonstrated that direct interaction with real-life agents draws even more attention and involves different networks across prefrontal cortical and amygdala areas than watching stills or even videos of the very same agents^10,22^. Neurons in the anterior cingulate cortex predict the behavior of others during social interaction in an iterated prisoner’s dilemma task and disruption selectively hampers mutually beneficial interactions in rhesus macaques^9^. A network in the dorsomedial prefrontal cortex codes for social partner identity and tracks these partners’ behavior as well as the distribution of food rewards in a real-life social task^23^. Thus, studies employing real-life interactions promise to shed new light on the processes involved in social information processing particularly if agents are freely moving^24^.

Social attention is dictated by the affordances of the stimulus situation, but can also vary considerably between individuals. Social status within a group can be a strong predictor of how much attention an individual pays their group mates^25–27^. In a population with clearly defined and repeatable inter-individual differences in sociability, low-sociable rhesus macaques paid more attention to any social information in videos than high-sociable monkeys independent of social status^28^. Early life adversity, such as experimental maternal immune activation during gestation, can also affect social attention in juvenile offspring possibly via epigenetic effects^29^. Both the genotype upstream of the serotonin transporter gene^30,31^ and administered serotonin precursor^32^ affect social attention, suggesting the involvement of a serotonergic pathway in generating the observed inter-individual differences^33^. Additionally, an oxytocinergic pathway has been implicated in linking individual differences in affiliation and dominance with visual attention^34^.

This study focuses on how individuals selectively attend to 3^rd^ person situations, i.e. interactions among others^13^, as they spontaneously unfold (or not) in the vicinity, excluding interactions involving themselves. We propose that the relevance of agonistic and affiliative 3^rd^ person interactions differs considerably, and that relational aspects add an additional layer of relevance^35^. For example, when conflicts arise, they can quickly escalate and involve the subject directly, either through redirected aggression^36^ or by soliciting their help^37^. As a result, social attention for aggressive interactions may serve a risk avoidance function and is likely modulated by the dominance relationship between the subject and the interacting parties. In contrast, the relevance of affiliation between others is often less immediate and typically indirect. Rates of grooming and agonistic support are often correlated^37^ which means that the state of the grooming network provides indirect information about the alliance network in the group^38^. Information about 3^rd^ person grooming will affect the subject only indirectly and usually not immediately, but when it needs to decide in a later situation whom to support or which affiliative relationships to invest in. Yet sometimes, primates also directly and immediately interfere in the grooming interactions of others to gain access to desirable partners or to manipulate others’ relationships^39,40^.

In a population of wild Assamese macaques in Thailand, we used behavioral event recording to quantify the overt attention of 56 adult males to 962 agonistic and affiliative 3^rd^ person interaction unfolding in subject’s vicinity (≤5m) or controls where two other adult males came into close proximity of ≤1.5m without interacting by means of vocalization, facial expression, touch or pose gesture. With statistical models we show that valence of the social scene (agonism, affiliation, control) interacted with dominance relations and social bond strength between the subject and the two stimulus monkeys to predict the occurrence of overt attention to the scene suggesting non-additive integration of different relevancies during processing of 3^rd^ person real-life interactions. We further identify individual differences in sustained attention for agonistic or affiliative interactions that are independent of subject rank. Note that we refer to the affiliative relationship between two males as their social bond strength to enhance accessibility of the results; strictly speaking, social bonds are only the strong, equitable, stable (and supportive) among all affiliative relationships^41^.

## METHODS

### Study site and subjects

This study was conducted at the Phu Khieo Wildlife Sanctuary in North-Eastern Thailand (PKWS, 16^⍰^ 05’-35’ N, 101^⍰^ 20’-55’ E, 1573km^2^) as part of a long-term research project on a wild population of Assamese macaques ongoing since 2005^42^. PKWS is part of the well-preserved Western Isaan Forest Complex covering an area of more than 6,500km^2 43^, which houses a diverse community of large mammals and predators, an indicator of low disturbance levels^43^. We collected data on 56 adult Assamese macaque males belonging to five different neighboring multimale–multifemale groups with 39 to 75 individuals each and a minimum of eight adult males and ten adult females.

### Social attention data collection

Social attention data were collected from June 2022 until April 2023 through event sampling of behavior towards social scenes spontaneously unfolding in the group. A social scene was considered valid for recording each time a stationary adult male subject in the vicinity (≤ 5 m) had an unobstructed view and the human observer clearly saw the onset of the social scene. We recorded head gaze (but also eye gaze if possible) towards three different social scene *conditions* that all started with two adult males (the stimulus individuals) arriving within 1.5 m of proximity of one another and continued either (1) as non-interacting *controls* if the males remained in close proximity without explicitly interacting via touch, vocalization or facial expression or (2) as *affiliation* if the two males only exchanged friendly behaviors, and (3) as *agonism* if the interaction included aggressive or submissive behaviors. Given the specificity of parameters and viewing conditions necessary to accurately record each attention event, these data were collected opportunistically outside the focal animal follows. Upon the unambiguous onset of a social attention event, we recorded the following variables: identities of the subject and two interacting stimulus individuals, behaviors displayed during the stimulus interactions, start and end times of stimulus interactions, timing of looks relative to the onset of the stimulus interaction and duration of each look. We additionally scored the interaction for conspicuousness (0/1) as a means to control statistically for loud or tumultuous interactions drawing attention^35^. We collected data from 962 social attention events, i.e. 17±9 events for each of the 56 subjects.

### Collection of social relationship data

Concomitant with the social attention data collection, all adult males were systematically observed as focal animals in 40 min protocols^44^ from June 2022 through April 2023. We recorded continuously ^45^ all affiliative and agonistic (both aggressive and submissive) interactions including the identities of the actor and receiver as well as all approaches into and departures from a 1.5 m sphere around the focal animal. To inform our dominance and social bond metrics (see below) about the state of relationships at the onset of the study in June 2022, we included data from January-May 2022. A total of 1,915.9h of continuous focal animal data were collected (mean ± standard deviation per male 33.6h ± 12.1).

### Dominance scoring – Elo-rating

For the assessment of dominance, we scored submissive behaviors (i.e. bare teeth, make room, give ground) in decided dyadic conflicts (either unprovoked submission or submission following aggression) registered during our continuous focal animal observations and ad libitum recordings from January 2022 through April 2023. To avoid long burn-in periods when males immigrate or mature, we employed Bayesian Elo-rating^46^ that estimates start values from the data and captures the state and dynamics of male dominance relationships, which can fluctuate remarkably in Assamese macaques^47^. We used the elo_seq_bayes and extract_elo_b functions of the EloRating.Bayes package^48^ in R version 4.3.0^49^. A short burn-in period of one month was applied for each male immigrating into a study group or maturing into an adult male based on his morphological characteristics (fully developed testes and canines, full body length larger than that of an adult female). For our statistical analysis, we extracted Elo-scores for the day that attention to a social scene was observed, modeled Elo-scores standardized per group^50^ and include group ID as a random effect.

### Social bond strength – DDSI

As a measure of the strength of dyadic social bond strength, we calculated Dynamic Dyadic Sociality Indices (DDSI) and followed De Moor et al.^51^ and Stranks et al.^47^ in their application of this method originally developed by Kulik and Mundry^52^. We focused strictly on adult male dyads, including only interactions between adult males in our computations. We based our DDSI scores on the time each dyad spent in body contact and allo-grooming, with each of these behaviors being weighed by the inverse of their frequency. Scores for all dyads started at the standard value of 0.5 (DDSI scores range from 0 to 1), indicative of a neutral relationship. Since affiliative interactions were rare, DDSI values were not well differentiated during the first half of the study. Thus, in all models, social bond strength was estimated as the DDSI value for that dyad at the end of the study period when calculations included a maximum of information.

### Statistical analysis

#### Looking probability model

We first tested the probability of a subject to look versus to not look at a scene as the response variable. Looks were only considered when occurring in the first 5 seconds after the start of social scene or else we scored a zero. To estimate how the probability of looking at a particular social scene varies across different conditions, and combinations of dominance relation and social bond strength, we fitted a Generalized Linear Mixed Model (GLMM)^53^ with binomial error structure and logit link function^54^. We included a three-way interaction term between condition (agonism, affiliation, and non-interacting control), dominance relation between the subject and the higher ranking of the two stimulus individuals (*ΔR_sub-max_* = Elo subject minus Elo higher ranking stimulus), and maximum DDSI value between the observer and each one of the interacting partners (*DDSI*_*sub-max*_). After we found that the three-way interaction term was not significant (see supplementary materials) we excluded it and ran a reduced model including the two-way interactions between the terms of the previous three-way interactions (see Table S2). In a last step, we excluded the non-significant interaction term dominance relation x max. DDSI and report below the final model including only two of the three 2-way interaction terms. In all models, we included rank of the subject (*R_sub_*) as well as conspicuity of the interaction as further fixed effects.

To control for their effects and avoid pseudo-replication, we included random intercept effects for study group, identity of the subject, identity of the actor, and identity of the receiver in the social scene. To avoid an overconfident model and secure a type I error rate at the nominal level of 0.05 we included all theoretically identifiable random slopes^55^, namely those of the three-way interaction term and all fixed effects within each random intercept. Though we initially aimed at including also all correlations among random intercepts and slopes^56^, we were forced to exclude them to allow for successful model convergence.

Prior to fitting the model, we first re-leveled the different levels of our categorical predictor for condition, defining controls (non-interacting) as the reference level for more intuitive result interpretation. We then inspected the distributions of dominance relation (*ΔR_sub-max_*), maximum DDSI (*DDSI_sub-max_*), and rank of the subject (*R_sub_*), which overall appeared roughly symmetrical and free of outliers. We z-transformed these predictors to a mean of zero and standard deviation of one to ease model convergence. We manually dummy coded and centered the control predictor (conspicuity of interaction) before including it as a random slope. The null model comprised only conspicuousness and the random effects.

The sample analyzed with the looking probability model comprised a total of 962 social attention situations, of which 337 were non-interacting controls, 354 affiliative interactions, and 271 agonistic events. These observations concerned a total of 56 subjects and 57 stimulus individuals. Looks occurred in a total of 364 situations, out of the total 962.

#### Looking duration model

In order to quantify variation in the duration of sustained attention, we ran a looking duration model using a subset of our full dataset corresponding only to situations where looks occurred. Here, we used *looking time* as a response, which had been widely used in previous human and non-human primate studies as an indicator of preferences and discriminative abilities^57^, potentially indicating a certain type of motivation to sustain attention.

To estimate the extent to which looking time varied across different conditions, and combinations of dominance relations and social bond strengths, we fitted a Generalized Linear Mixed Model (GLMM)^53^ with gamma error structure and log link function^58^. This model was strictly identical to our looking probability model in both fixed and random effects structure. Nonetheless, to control for the effect that longer social scenes may have on looking duration, we here included the duration of the stimulus interaction as a further fixed effect. We removed all correlations among random intercepts and slopes^56^ in an attempt to achieve successful model convergence. Since we faced convergence issues from the start, we ultimately fitted a Bayesian model using the brms package in R. The model was run using 4 MCMC chains of 4,000 iterations per chain with adapt_delta set at 0.99 and a max_treedepth of 15, including 1,000 “warm up” iterations during which no divergent transitions were reported. This resulted in 12,000 posterior samples per model.

We assessed the distributions of dominance relations (*ΔR_sub-max_*), maximum social bond strength (*DDSI_sub-max_*), and rank of the subject (*R_sub_*), which overall appeared roughly symmetrical and free of outliers. We z-transformed these predictors to a mean of zero and standard deviation of one to ease model convergence. We further inspected the distribution of the response (looking duration) and the added fixed effects of duration of the stimulus situation, both of which had heavily right-skewed distributions. To deal with this, we log transformed interaction duration. Given that we were fitting a model with a log link function, we kept the response as is. We manually dummy coded and centered the control predictor (conspicuity of interaction) before including it as a random slope. We defined weakly informative priors for all model parameters as to not strongly restrict posterior distributions. For all fixed effects we used a Normal (0, 1) prior, Exponential (1) priors for the standard deviations of random effects, and Exponential (1) for the shape parameter.

The sample analyzed with this model comprised a total of 441 social scenes that were looked at including those where looks occurred later than 5 seconds after scene onset. Ninety-nine scenes were non-interacting controls, 160 affiliative interactions, and 182 agonistic interactions. These observations involved 56 different males as subjects. Looking duration ranged from 1 to 49 seconds.

#### Statistical model implementation

The looking probability model was fitted in R (version 4.4.3)^49^, using the function glmer of the package lme4 (version 1.1-35.1)^59^. We determined confidence limits of model estimates and fitted values by means of a parametric bootstrap (N=1,000 bootstraps; function bootMer of the package lme4). We determined model stability by dropping the levels of the grouping factors one by one, fitting the full model to each of the subsets, and finally comparing the range of estimates with those obtained for the full data set. This revealed moderate to good stability of the estimates (see Table 1 in the results section). We determined the significance of individual fixed effects by means of likelihood ratio tests^60^ comparing the full model with models lacking them one at a time (R function drop1).

**Table 1.**
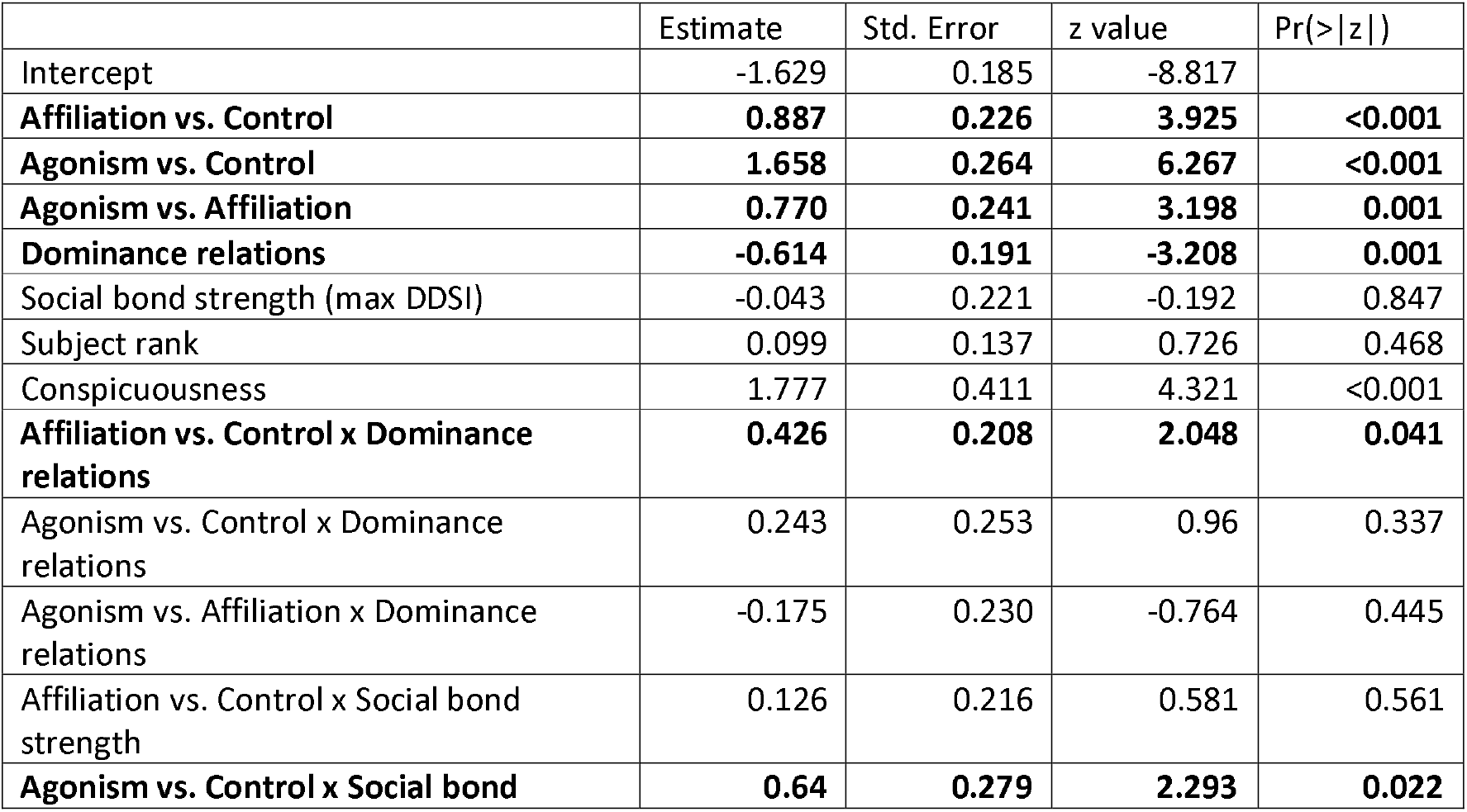
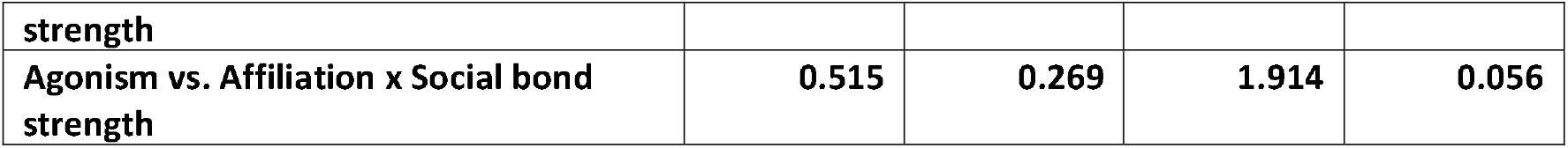
Model estimates for the reduced looking probability model, including as predictors only the interaction term between condition and dominance relations between the subject and the dominant of the stimulus individuals. The full model was significantly different from the null model including only control factors (conspicuousness of the interaction), and random effects (Chi^2^ = 34.02, df = 11, P<0.001). Significant effects of test predictors are highlighted in bold. Significance of the different levels of condition was assessed by releveling the intercept.

The looking duration model was fitted in R (version 4.4.3)^49^, using the function brm of the package brms (version 2.22.0), which builds on the computational framework Stan [https://mc-stan.org] to fit Bayesian models^61^. The convergence and stability of the models were assessed with R-hat values, which were below 1.01^62^ and a visual inspection of the plots of the chains^63^. In addition, all Bulk and Tail Effective Sample Sizes (ESS) for the estimated posteriors were >6000 ^61,62^.

## RESULTS

Results are based on the behavior of 56 male subjects that had the opportunity to visually attend to 962 social scenes involving any combination of two males co-residing in their group. Social scenes included 271 agonistic events, 354 affiliative interactions, and 337 non-interacting control situations where two adult males were in close proximity of each other and did not interact via vocal exchange, facial expression, or tactile behavior.

### Selective attention (looking probability)

In the looking probability model, the three-way interaction term between condition, dominance relations, and maximum DDSI was not significant and subsequently was excluded from our first reduced model (Tab. S1). To arrive at our final model, we further excluded the non-significant two-way interaction maximum DDSI x dominance relations (Tab. S2). The final full model differed significantly from the null model including only the control variable conspicuity of the interaction and the random effects of subject ID and group (Chi^2^ = 34.02, df = 11, P<0.001).

Based on the parameter estimates and conditional on the interaction effects, subjects were more likely to attend to interactions than to non-interactive controls (Tab. 1, Fig. 2). The higher above the subject the higher-ranking of the stimulus individual was, i.e. the more negative the dominance relations, the more likely was the subject to pay attention to an agonistic interaction and the lower below the subject the stimulus individuals were, the less likely the event was attended to. This effect of dominance relations on overt attention differed between conditions. Specifically, the effect of dominance relations was stronger (more negative estimate) for non-interacting controls than for affiliative interaction stimuli, but not stronger than for agonistic scenes.

**Figure 1.**
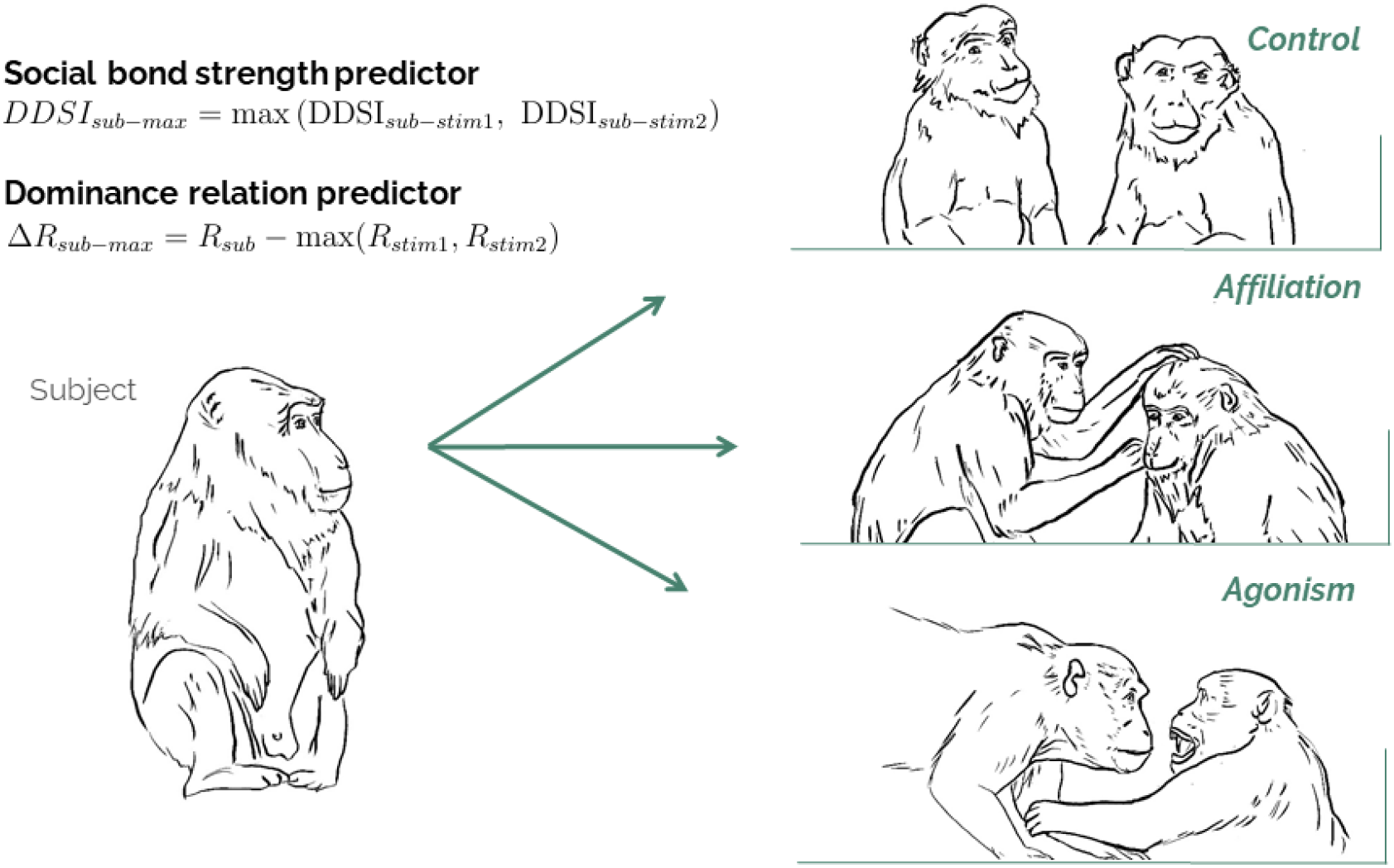
This field study integrates data on visual attention to social scenes spontaneously unfolding in the vicinity of a subject (sub) with information about the affiliative and dominance relationships among all males in five study groups. Social scenes with two adult males in close proximity were classified as non-interacting controls or interactions reflecting affiliation or agonism. The relationship of the subject to the two stimulus males (stim 1, stim 2) in the scene was quantified as the dominance relation to the higher-ranking of the stimulus males (Rank difference sub-max) and the maximum social bond strength to either of the two (DDSI_sub-max_).

**Figure 2.**
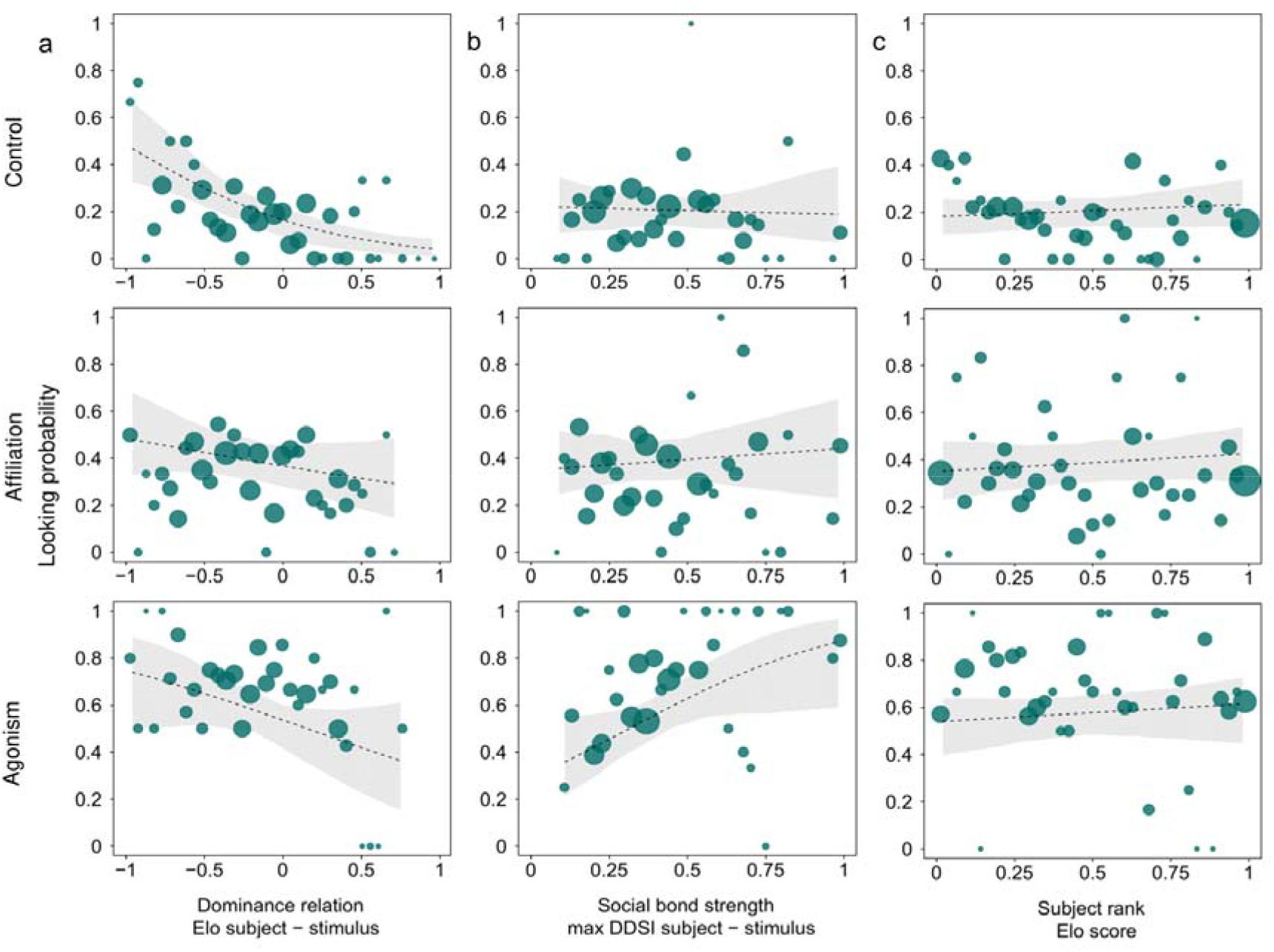
Variation in visual attention to 3^rd^ person social scenes as predicted from the looking probability model. Subjects generally attended more often to interactions than control scenes and to agonistic than affiliative interactions. Both the 2a) negative effect of dominance relations (subject minus dominant stimulus) and the 2b) positive effect of social bond strength (subject to stimuli) were modulated independently by condition and were stronger for agonistic than for affiliative interactions. Subject rank 2c) did not affect visual attention. Bubbles are average probabilities per observed value of the predictor with size representing the number of observations, lines are model fitted estimates and shades their confident intervals (Tab. 1). Data came from 56 subjects and 962 social scenes.

Furthermore, stimulus valence modulated the effect of social bond strength on visual attention. The closer the subject was affiliated to the stimuli (the higher the maximum DDSI), the more likely they paid attention which was an effect exacerbated for agonistic interactions, still pronounced for affiliative 3^rd^ person interactions and weakest for non-interacting control scenes.

Beyond the relational effects of dominance relations, the individual dominance rank of the subject did not affect its probability to monitor stimuli across conditions (see Fig. 3). Situations that were more conspicuous were more likely to be looked at.

**Figure 3.**
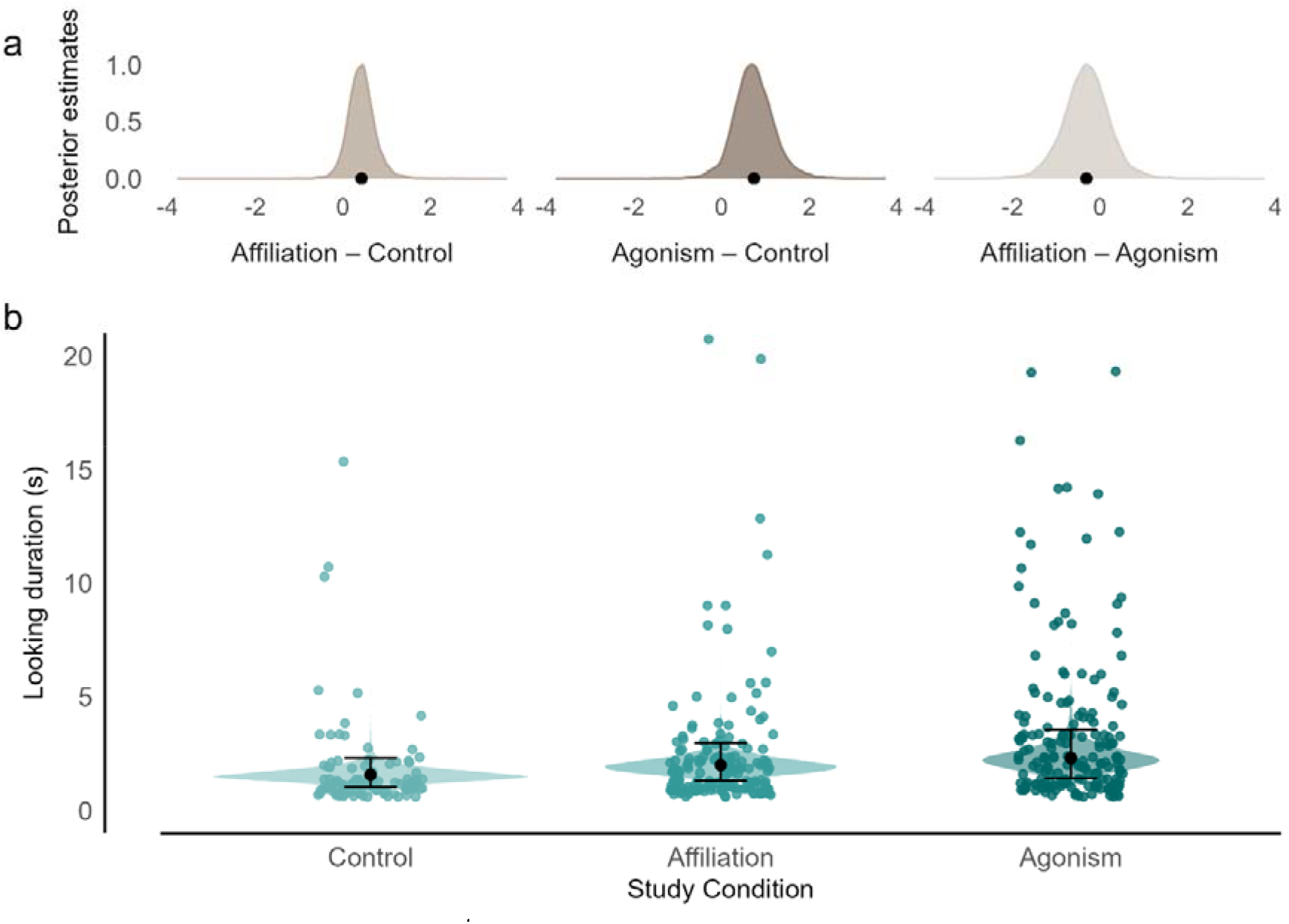
Sustained attention for 3^rd^ person interactions was longer than for non-interaction social control scenes and longer for agonistic than for affiliative interactions (Tab. 2) with distributions of posterior estimates for the looking duration model shown in 3a). Violin plots are drawn for raw data 3b) and whisker plot shows model estimates with 95% credible interval.

**Table 2.** Model estimates for the reduced looking duration model including only main effects; posterior means are reported with their standard errors and 95% credible intervals for the fixed effects. Estimates are based on 4,000 posterior draws.

|  | Estimate | Est.Error | Q2.5 | Q97.5 |
| --- | --- | --- | --- | --- |
| Intercept | 0.171 | 0.206 | -0.234 | 0.573 |
| **Affiliation vs. Control** | **0.239** | **0.164** | **-0.086** | **0.565** |
| **Agonism vs. Control** | **0.380** | **0.215** | **-0.082** | **0.774** |
| Agonism vs. Affiliation | 0.141 | 0.244 | -0.362 | 0.592 |
| Dominance relations | 0.014 | 0.105 | -0.198 | 0.218 |
| Social bond strength (max DDSI subject – stimulus) | 0.035 | 0.109 | -0.184 | 0.251 |
| Subject rank | -0.042 | 0.121 | -0.263 | 0.194 |
| Log interaction time | 0.095 | 0.058 | -0.016 | 0.210 |
| Conspicuousness | 0.359 | 0.178 | -0.020 | 0.684 |

### Sustained attention (looking duration)

We used a Bayesian model to estimate the effects of the same predictor variables on the time spent looking at the social scene. Posterior probabilities for the three-way-interaction between dominance relations, social bond strength and condition were centered around zero (Tab. S3). In a reduced model without the 3-way interaction term, we found no evidence supporting either of the three 2-way interaction effects (all posterior probabilities centered around zero). The final model (Tab. 2) including only main effects provided clear evidence for sustained attention to vary with condition. Males demonstrated longer attention for agonistic interactions between other males than for scenes with two non-interacting stimulus males (posterior probabilities >0.95 for difference between control and each of agonistic and affiliative scenes; Fig. 3). Furthermore, there was moderate evidence for longer looking time towards agonistic compared to affiliative interactions (posterior probability 0.84). Looking time during agonistic situations was on average 0.55s longer compared to non-interacting controls, and 0.23s longer than affiliation. Dominance relations and social bonds to the stimulus males and subject rank did not affect sustained attention.

### Inter-individual differences

Random effects from both models suggest that individuals do not systematically differ in attention to just any type of social scene. Instead, significant individual differences were specific to and differed between sustained attention to affiliative scenes (mean individual random effect for affiliative scenes ± stddev. = 0.0013 ± 0. 174; Fig. 4a) and agonistic scenes (0.0012 ± 0.192; Fig.4b). These random effects were extracted from a model with and are therefore controlled for subject dominance rank.

**Figure 4.**
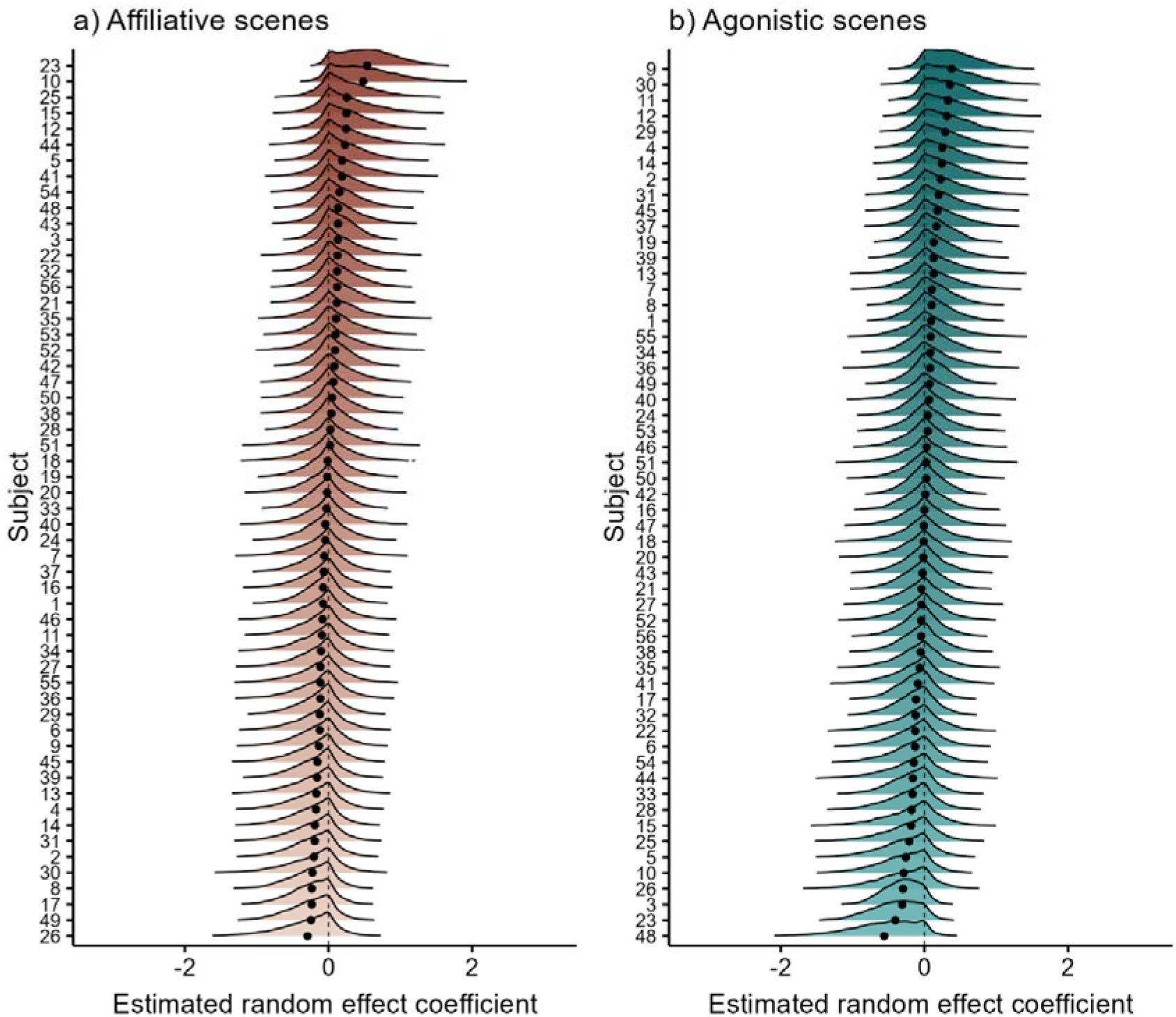
Inter-individual variation in average looking duration. Posterior distributions are shown for a) affiliative scenes and b) agonistic scenes for all study subjects relative to the estimated population average. Black dots show the posterior mean for each subject. Positive estimates indicate above-average looking durations, whereas negative estimates indicate below-average looking durations.

## DISCUSSION

Our study contributes to the development of real-life social neuroscience by elucidating the neural mechanisms underlying social information integration. Social interactions are inherently complex, requiring individuals to make predictions not only about their own actions but also about those of others, informed by observations of 3^rd^ person interactions^13^. We found that individuals allocate increased visual attention to 3^rd^ person interactions, particularly those involving agonistic versus affiliative behaviors, and that different dimensions of social relationships between the subject and third parties independently modulate this attentional bias. Furthermore, we observed significant individual differences in sustained attention to others’ interactions, underscoring the importance of considering personal factors in understanding social cognition. These results have important implications for our understanding of how the brain integrates social information and how emotion regulation and adaptive benefits influence sustained attention.

### 3^rd^ person interactions draw more attention than non-interacting social controls

Using as a control a social scene with two individuals in close proximity without an interaction, we clearly demonstrated that both the probability of overt visual attention and the duration of sustained visual attention were increased for 3^rd^ person interactions. The three social conditions were matched for salience by requiring the scene to have a clear starting point where two males arrived in close proximity, i.e. all scenes were initially dynamic. By requiring this conservative control condition, we exclude the possibility that interaction scenes draw more attention, simply because perceptually two individuals may be more salient^21^. Exceedingly dynamic scenes that involved quick movements and/or vocalizations either in the agonistic or the affiliative context (e.g. ritualized male-infant-male interactions)^64^ were scored to be conspicuous which was a control variable in all models. Thus, visual attention was not only guided by stimulus salience alone, but also by the mental representation of its social relevance^6,65,66^ and was sustained into a range where MEG studies suggest different information integration events can be detected (1,100 and 2,000 ms)^67^. To our knowledge, the only other study to demonstrate that 3^rd^ person interactions attract more visual attention is a three player experiment with rhesus macaques that identified neurons in the prefrontal cortex to be differentially activated when two others interacted^23^.

### Attention guided by valence of others’ interactions

Visual attention was further guided by interaction valence, adding ecological validity to previous findings on selective attention for and differential processing of threat faces over photos of friendly teeth-chattering^2,68^. Here the naturalism scale seems particularly relevant, because the two facial expressions differ in important ways; teeth chattering is a repetitive and thus inherently more dynamic and auditory display than a threat face and therefore may be more difficult to identify from a still photograph. Even if videos of agonistic versus affiliative interactions of conspecifics are used, chimpanzees attend equally long to both conditions^21^. Yet even in our real-life scenes in proximity of the subject, monkeys attended more to agonistic than affiliative interactions of others.

Agonistic interactions may be more relevant, because the subject could be drawn into the conflict by redirected aggression^69^ or be solicited to help one of the opponents^70,71^. The fact that watching others fight increases rates of self-directed behaviors indicative of increased emotional arousal^72,73^, suggests that subjects do not only gather information about the state of the dominance relations of others, but also about imminent threat to themselves. Others’ affiliative interactions may also lead to direct involvement of the subject, if they decide to interfere by joining or stopping the affiliation^39,40^, but this decision lies with the subject and not with a 3^rd^ party and may therefore not require the same level of attention as 3^rd^ person agonism.

Despite the primacy of agonism, subjects still paid more often and more sustained attention to 3^rd^ person friendly interactions than non-interacting controls. Primates look towards the mother (the main affiliate), if they hear an infant in distress from the other direction^74^ and look longer at a staged conflict, if it is between close affiliation partners compared to other dyads^75^. Selective attention for 3^rd^ person friendly interactions must underlie such findings, but had not been previously demonstrated.

### Two dimension of social relationships independently guide visual attention

Variation in imminent threat may also explain how the social relationships between the subject and the stimulus individuals have moderated valence effects on selective attention. Since aggression flows down the hierarchy in macaques^76^, imminent threat increases the higher above the subject the dominant stimulus male. This effect appears most pronounced in non-interactive control scenes, because in controls, attention drops to zero for scenes that only have males subordinate to the subject; at the largest dominance differences (left of Fig. 2a) control scenes did not attract more attention than agonistic or affiliative scenes. Attention to non-interacting control scenes with individuals out-ranking the subject may serve in uncertainty-reduction^77^. Attention for conflicts was much higher than for controls, even when both males were subordinate to the subject, so that the rank-difference had a slightly weaker effect. Dominance difference did not moderate visual attention to affiliative interaction scenes which suggests that it is not the mere presence of a higher ranking individual that adds relevancy, but the action of a dominant.

We expected that monitoring close affiliates may also contribute towards risk-avoidance, given that the conflicts of closely bonded partners are the most likely to trigger risky interactions for the observing party^78,79^ which requires behavioral coordination between close affiliates^36,80^. Matching these expectations, we found the attention-grabbing effect of social bond strength with stimuli to vary with stimulus valence: the social bond effect was most pronounced for conflicts and less so for affiliation and control scenes. This highlights again the increased relevance of potentially risky situations. In other real-life observational studies on captive populations, female rhesus macaques and mandrills of both sexes also paid more attention to their closest partner and their kin, respectively^35,81^. This is in contrast to a recent study on male rhesus macaques^34^; the grooming relationship did not affect visual attention for pictures of a group mate, whereas the aggressive relationship with a stimulus clearly correlated with attention for this male^34^. By moving from the presentation of still photographs to real-life stimuli and from individual stimuli^81^ to interactions, we demonstrate that two dimensions of a subject’s relationships to others have independent effects on the allocation of visual attention which is more pronounced for agonism than affiliation of others.

### Individual differences in social attention

For good reasons, the most controlled studies employing brain-imaging or electrophysiology often suffer from small sample sizes which unfortunately prohibits investigation of individual differences in social attention and underlying processing^1^. Here, we find no evidence for individuals to generally differ in their social attention or for social status. Instead, the 56 male Assamese macaques varied in different ways in how they attended to agonistic interactions or to affiliation between others.

In a well-powered study on rhesus macaques, significant year-to-year repeatability was found in attention to threat over neutral faces, but not in attention to friendly teeth-chattering over neutral faces^82^. Differences between four rhesus macaques in the propensity to give aggression, grooming, and approaches over receiving these behaviors were all associated to some degree with the social attention they gained in a distractor task^34^. Neither of the rhesus macaque studies, however, controlled for dominance rank prior to extracting information on individual differences. We can clearly show here that despite dominance changing throughout the study period, individuals maintain differences in how they attend to conflicts of others, suggesting stable differences in underlying regulation via serotoninergic and oxytocinergic pathways^33,34^.

### Conclusion – information integration for visual attention

Together, these results suggest that different relevancies of social information are integrated in complex ways. Interaction versus non-interacting control, valence of the interaction, and two separate relational aspects do not have simple additive effects on social attention. The social value of attention is evident from increased attention to others’ affiliation over non-interacting controls that is unaffected by dominance differences or social bonds towards the stimuli. All other results can be explained by risk-avoidance alone where information is not gathered for its own sake, but emotion regulation prepares and activates the subject in the face of imminent danger. Thus, social attention serves at least two goals in tandem, which explains why it involves several brain areas in the dorsomedial prefrontal cortex, dorsal anterior cingulate cortex, the ventral striatum and the amygdala perhaps with pronounced mixed selectivity at the neuronal and network level^9,23,83,84^.

## Supporting information

Supplementary results

## ACKNOWLEDGEMENTS

We are grateful to the National Research Council of Thailand and the Department of National Parks, Wildlife and Plant Conservation of Thailand for permission from to conduct research in a protected area (permits: 0907.4/9313 April 2^nd^ 2022, 0907.4/19458 September 3^rd^ 2023). We thank the superintendent of PKWS Wichanon Saenphala for his collaboration, Roger Mundry for statistical advice, and the team of foreign and Thai field researchers that supported data collection. This study was funded by the Deutsche Forschungsgesellschaft DFG-SFB 1528 Cognition of Interaction (project-ID 454648639).

## AUTHOR CONTRIBUTIONS

SP Investigation, Data curation, Formal analysis, Visualization, Writing - Original draft

SMa Resources, Project administration, Writing - Review & editing

SMe Resources, Project administration, Writing - Review & editing

JO Conceptualization, Funding acquisition, Data curation, Resources, Supervision, Writing - Review & editing

OS Conceptualization, Funding acquisition, Data curation, Resources, Supervision, Writing - Original draft

## CONFLITCS OF INTEREST

The authors declare no conflict of interest.

## DATA AVAILABILITY

Replication data are available at GRO.data https://doi.org/10.25625/XFN4B4

## ETHICAL STATEMENT

The study was conducted with permission of relevant Thai authorities under a benefit sharing agreement and was purely observational, completely non-invasive and did not involve food provisioning or capture of the animals.

