## Supplementary results for "Complex modulation of visual attention to 3^rd^ person interactions in wild macaques"

for

**Table S1.** Model estimates for the **full** **looking probability model**, including as predictors a three-way interaction term between condition, dominance relation between the subject and the dominant of the stimulus individuals, and maximum social bond strength (DDSI) between the subject and the stimulus dyad. This full model was significantly different from the null model including only control factors (conspicuity of the interaction), and random effects (Chi^2^ = 34.02, df = 11, P<0.001). Significant fixed effects are highlighted in bold. Significance of the different levels of condition was assessed by releveling the intercept.

|  | Estimate | Std. Error | z value | Pr(>\|z\|) |
| --- | --- | --- | --- | --- |
| Intercept | -1.659 | 0.191 | -8.698 |  |
| **Affiliation vs Control** | **0.916** | **0.248** | **3.693** | **<0.001** |
| **Agonism vs Control** | **1.664** | **0.264** | **6.311** | **<0.001** |
| **Agonism vs Affiliation** | **0.763** | **0.240** | **3.186** | **0.001** |
| **Dominance relations** | **-0.589** | **0.195** | **-3.012** | **0.003** |
| Social bond strength (max DDSI) | 0.005 | 0.222 | 0.024 | 0.981 |
| Rank subject | 0.093 | 0.135 | 0.687 | 0.492 |
| **Conspicuousness** | **1.77** | **0.404** | **4.382** | **<0.001** |
| **Affiliation vs Control x Dominance relations** | **0.408** | **0.214** | **1.904** | **0.057** |
| Agonism vs Control x Dominance relations | 0.199 | 0.261 | 0.762 | 0.446 |
| Agonism vs Affiliation x Dominance relations | -0.179 | 0.234 | -0.765 | 0.444 |
| Affiliation vs Control x Social bond strength | 0.048 | 0.228 | 0.208 | 0.835 |
| **Agonism vs Control x Social bond strength** | **0.568** | **0.277** | **2.054** | **0.040** |
| **Agonism vs Affiliation x Social bond strength** | **0.520** | **0.274** | **1.899** | **0.058** |
| **Dominance relations x Social bond strength** | **0.325** | **0.180** | **1.800** | **0.072** |
| Affiliation x Dominance relations x Max final DDSI | -0.357 | 0.240 | -1.492 | 0.136 |
| Agonism x Dominance relations x Max final DDSI | -0.436 | 0.267 | -1.631 | 0.103 |

**Table S2.** Model estimates for the **first reduced** **looking probability model**, including as predictors all possible two-way interaction terms between condition, dominance relation between the subject and the dominant of the stimulus individuals, and maximum social bond strength (DDSI) between the subject and the stimulus dyad. The full model was significantly different from the null model including only control factors (conspicuity of the interaction), and random effects (Chi^2^ = 34.02, df = 11, P<0.001). Significant fixed effects are highlighted in bold. Significance of the different levels of condition was assessed by releveling the intercept.

|  | Estimate | Std. Error | z value | Pr(>\|z\|) |
| --- | --- | --- | --- | --- |
| Intercept | -1.638 | 0.186 | -8.788 |  |
| **Affiliation vs Control** | **0.899** | **0.235** | **3.827** | **<0.001** |
| **Agonism vs Control** | **1.657** | **0.266** | **6.219** | **<0.001** |
| **Agonism vs Affiliation** | **0.750** | **0.281** | **2.670** | **0.008** |
| **Dominance relations** | **-0.609** | **0.193** | **-3.157** | **0.002** |
| Social bond strength (max DDSI) | -0.029 | 0.226 | -0.128 | 0.898 |
| Rank subject | 0.097 | 0.137 | 0.706 | 0.480 |
| **Conspicuousness** | **1.786** | **0.406** | **4.399** | **<0.001** |
| **Affiliation vs Control x Dominance relations** | **0.443** | **0.213** | **2.082** | **0.037** |
| Agonism vs Control x Dominance relations | 0.251 | 0.255 | 0.986 | 0.324 |
| Agonism vs Affiliation x Dominance relations | -0.179 | 0.240 | -0.745 | 0.457 |
| Affiliation vs Control x Social bond strength | 0.109 | 0.217 | 0.502 | 0.616 |
| **Agonism vs Control x Social bond strength** | **0.621** | **0.276** | **2.246** | **0.025** |
| **Agonism vs Affiliation x Social bond strength** | **0.542** | **0.277** | **1.955** | **0.051** |
| Dominance relations x Max final DDSI | 0.055 | 0.098 | 0.568 | 0.570 |

**Table S3.** Model estimates for the **full looking duration model**, including as predictors a three-way interaction term between condition, dominance relation between the subject and the dominant of the stimulus individuals, and maximum social bond strength (DDSI) between the subject and the stimulus dyad. The posterior means are reported with their standard errors and 95% credible intervals for the fixed effects. Estimates are based on 4,000 posterior draws.

|  | Estimate | Est.Error | Q2.5 | Q97.5 |
| --- | --- | --- | --- | --- |
| Intercept | 0.159 | 0.210 | -0.241 | 0.577 |
| **Affiliation vs. Control** | **0.246** | **0.174** | **-0.090** | **0.591** |
| **Agonism vs. Control** | **0.394** | **0.220** | **-0.067** | **0.800** |
| Agonism vs. Affiliation | 0.148 | 0.252 | -0.261 | 0.535 |
| Dominance relations | 0.003 | 0.128 | -0.246 | 0.251 |
| Social bond strength (max DDSI) | 0.020 | 0.133 | -0.245 | 0.273 |
| Subject rank | -0.039 | 0.120 | -0.268 | 0.206 |
| Log interaction time | 0.096 | 0.062 | -0.022 | 0.214 |
| **Conspicuousness** | **0.354** | **0.178** | **-0.024** | **0.682** |
| Affiliation x Dominance relations | 0.018 | 0.117 | -0.213 | 0.249 |
| Agonism x Dominance relations | 0.015 | 0.114 | -0.210 | 0.239 |
| Affiliation x Social bond strength | -0.015 | 0.112 | -0.239 | 0.202 |
| Agonism x Social bond strength | 0.046 | 0.111 | -0.174 | 0.261 |
| Dominance relations x Social bond strength | 0.065 | 0.092 | -0.116 | 0.245 |
| Affiliation x Dominance relations x Social bond strength | -0.092 | 0.113 | -0.315 | 0.131 |
| Agonism x Dominance relations x Social bond strength | -0.110 | 0.107 | -0.318 | 0.101 |

**Table S4.** Model estimates for the **first reduced looking duration model**, including as predictors all possible two-way interaction terms between condition, dominance relation between the subject and the dominant of the stimulus individuals, and maximum social bond strength (DDSI) between the subject and the stimulus dyad. The posterior means are reported with their standard errors and 95% credible intervals for the fixed effects. Estimates are based on 4,000 posterior draws.

|  | Estimate | Est.Error | Q2.5 | Q97.5 |
| --- | --- | --- | --- | --- |
| Intercept | 0.178 | 0.211 | -0.229 | 0.604 |
| **Affiliation vs. Control** | **0.236** | **0.170** | **-0.094** | **0.573** |
| **Agonism vs. Control** | **0.379** | **0.219** | **-0.073** | **0.783** |
| Agonism vs. Affiliation | 0.143 | 0.247 | -0.264 | 0.528 |
| Dominance relations | -0.018 | 0.126 | -0.265 | 0.226 |
| Social bond strength (max DDSI) | 0.020 | 0.135 | -0.244 | 0.284 |
| Rank subject | -0.043 | 0.121 | -0.268 | 0.199 |
| Log interaction time | 0.094 | 0.059 | -0.026 | 0.209 |
| **Conspicuousness** | **0.360** | **0.176** | **-0.011** | **0.678** |
| Affiliation x Dominance relations | 0.045 | 0.114 | -0.179 | 0.268 |
| Agonism x Dominance relations | 0.033 | 0.112 | -0.187 | 0.254 |
| Affiliation x Social bond strength | -0.009 | 0.113 | -0.233 | 0.208 |
| Agonism x Social bond strength | 0.048 | 0.113 | -0.175 | 0.267 |
| Dominance relations x Social bond strength | -0.017 | 0.048 | -0.111 | 0.078 |
